## Supplementary Information for "CRISPR targeting of H3K4me3 activates gene expression and unlocks centromere-proximal crossover recombination in Arabidopsis"

### SUPPLEMENTARY FIGURE LEGENDS:

S1

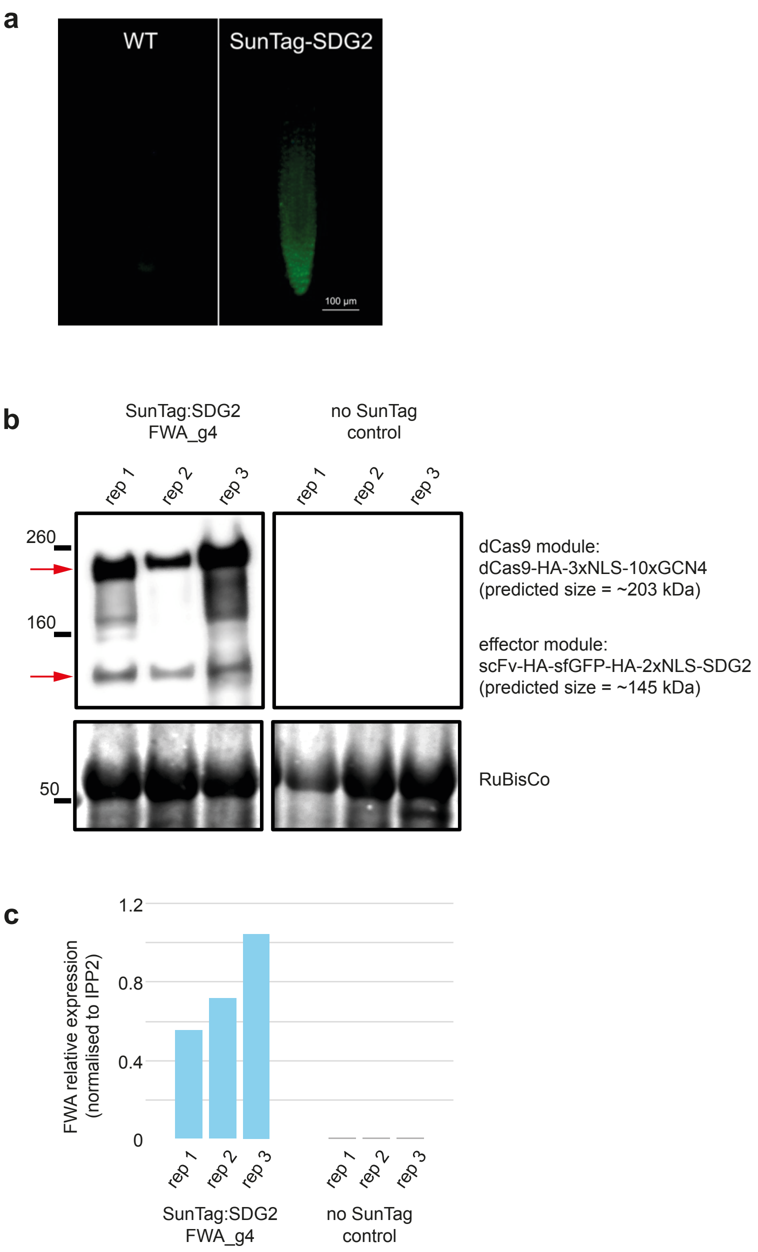

**Supplementary Figure 1:** A) epifluorescence image of rootstocks from 2-week old representative plants showing nuclear localisation of sfGFP from SunTag lines. B) Western blot for the dCas9-10xGCN4 and effector module in SunTag lines showing activation of FWA expression by RT-qPCR show in C). RT-qPCR for *FWA* relative expression (normalised to *IPP2*) from the same individual plants as used for western blot, shown in B).

S2

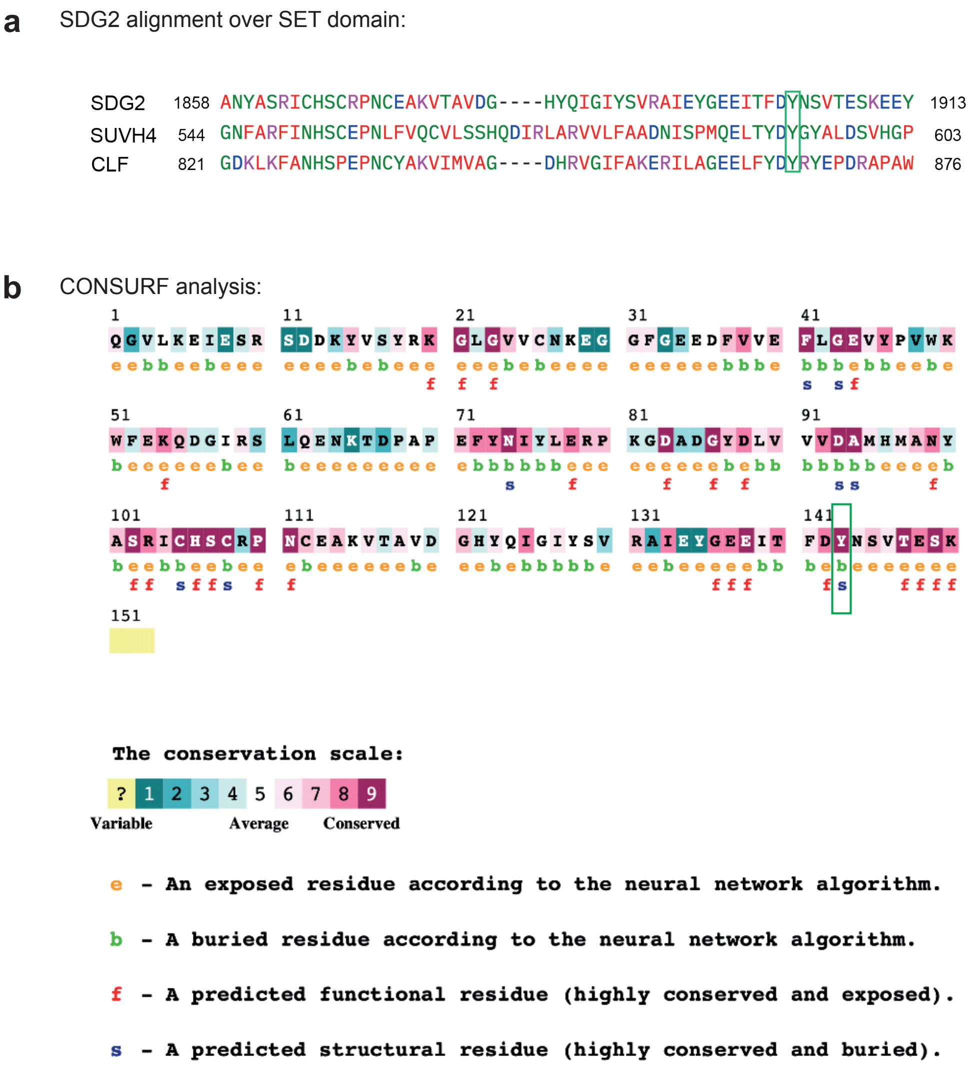

**Supplementary Figure 2:** A) Amino acid alignment between SDG2 and two other well characterised A. thaliana histone methyltransferases, SUVH4 (H3K9me2) and CLF (H3L27me3). The Y at amino acid position 1903 of SDG2 that was mutated to F in the SunTag:dSDG2 constructs corresponds to the Y593F in SUVH4(/KYP) that has previously been shown to abolish methyltransferase catalytic activity (Du et al., 2014) (position indicated in green rectangle). B) CONSURF analysis (Ashkenazy et al., 2016) on the SDG2 predicted SET domain (amino acids 1761-1911) showing the maximum level of conservation for SDG2$_{Y1903}$ (position indicated in green rectangle).

S3

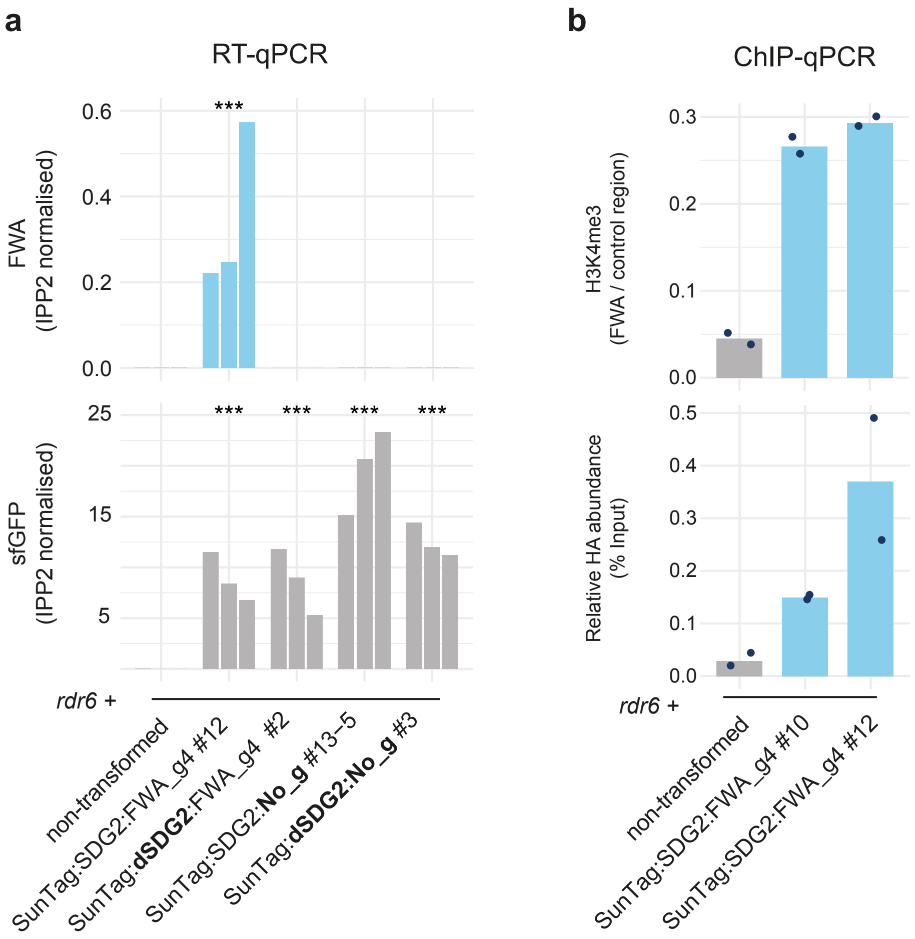

**Supplementary Figure 3:** A) qRT-PCR for the expression of FWA (upper panel) and sfGFP (lower panel) with individual biological replicates shown. *** indicates significant difference by ANOVA with post-hoc Tukey HSD for FWA, and Dunnett's for sfGFP (with non-transformed being control group), $p$ < 0.05. B) ChIP qPCR with two independent FWA targeting lines.

S4

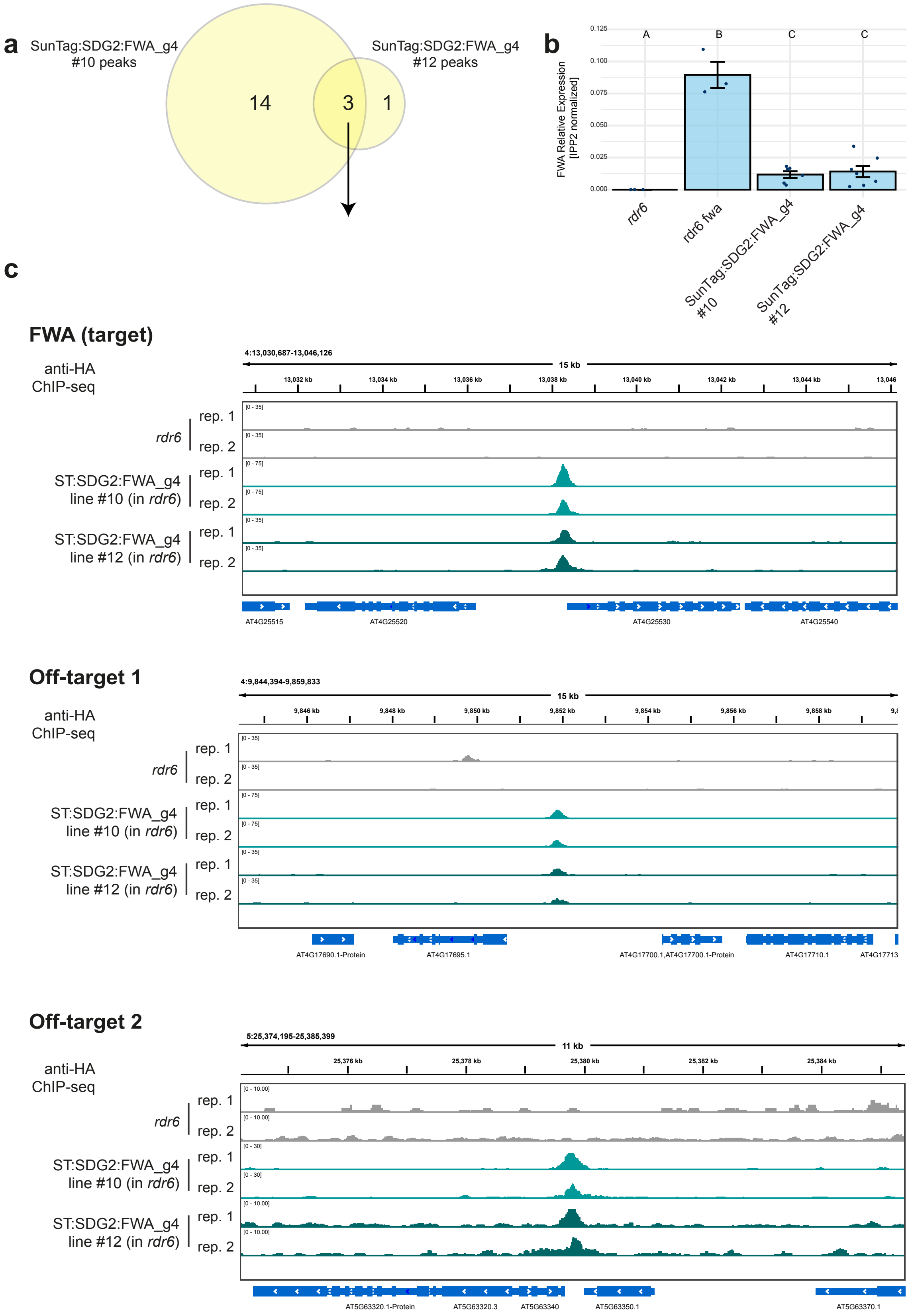
a
SunTag:SDG2:FWA_g4 #10 peaks
SunTag:SDG2:FWA_g4 #12 peaks
14
3
1
b
FWA Relative Expression [IPP2 normalized]
0.125
0.100
0.075
0.050
0.025
0.000
A
B
C
C
rdr6
rdr6 fwa
SunTag:SDG2:FWA_g4 #10
SunTag:SDG2:FWA_g4 #12
c
FWA (target)
anti-HA ChIP-seq
4:13,030,687-13,046,126
15 kb
rdr6
rep. 1
rep. 2
ST:SDG2:FWA_g4 line #10 (in rdr6)
ST:SDG2:FWA_g4 line #12 (in rdr6)
AT4G25515
AT4G25520
AT4G25530
AT4G25540
Off-target 1
4:9,844,394-9,859,833
AT4G17690.1-Protein
AT4G17695.1
AT4G17700.1,AT4G17700.1-Protein
AT4G17710.1
AT4G17713
Off-target 2
5:25,374,195-25,385,399
11 kb
AT5G63320.1-Protein
AT5G63320.3
AT5G63340
AT5G63350.1
AT5G63370.1

**Supplementary Figure 4:** A) Venn diagram showing the overlap between the peaks called from HA ChIP-seq of two independent SunTag:SDG2 FWA targeting lines. B) RT-qPCR analysis of FWA expression in two independent SunTag:SDG2:FWA_g4 lines. Dots represent individual biological replicates (a single plant) from the lines (#) and genotypes indicated. Error bars represent SEM. Different letters indicate significant difference by ANOVA with post-hoc Tukey HSD ($p$ < 0.05). n = 3 for *rdr6* and *rdr6 fwa,* n = 6 for SunTag:SDG2:FWA_g4 #10 and n = 7 for line #12. Note that while line #10 appears to bind more strongly to the FWA target than line #12, this does not correspond to a significant difference in FWA activation. C) Genome browser images for the overlapping peak regions, the top most significant peak is FWA, while the other two are off-target regions with sequence similarity to the FWA binding site that were previously identified (Papikian et al., 2019).

S5

a

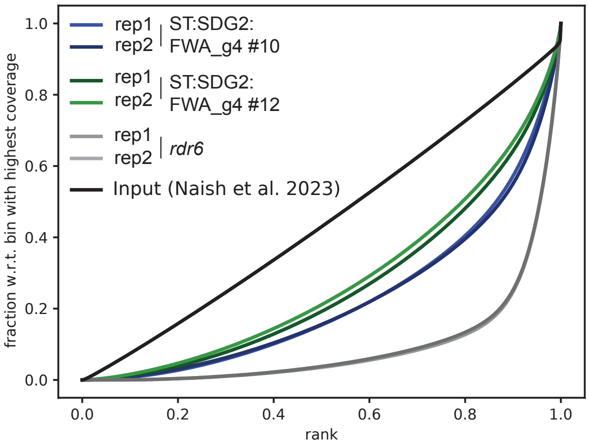

b

| Sample | Total mapped reads | mapped reads in peaks | FrIP (%) |
|---|---|---|---|
| SunTag:SDG2:FWA_g4 #10 rep 1 | 16218459 | 7845584 | 48.4 |
| SunTag:SDG2:FWA_g4 #10 rep 2 | 20650830 | 10570918 | 51.2 |
| SunTag:SDG2:FWA_g4 #12 rep 1 | 12360421 | 4619230 | 37.4 |
| SunTag:SDG2:FWA_g4 #12 rep 2 | 18237850 | 7013976 | 38.5 |
| rdr6 rep1 | 20389796 | 16361705 | 80.2 |
| rdr6 rep2 | 18015360 | 14238366 | 79.0 |

**Supplementary Figure 5:** Quality control for H3K4me3 ChIPseq of FWA targeting SunTag:SDG2 lines. A) Fingerprint plot (DeepTools) B) Table of total mapped reads and fragments of reads in peaks (FRiP) scores.

S6

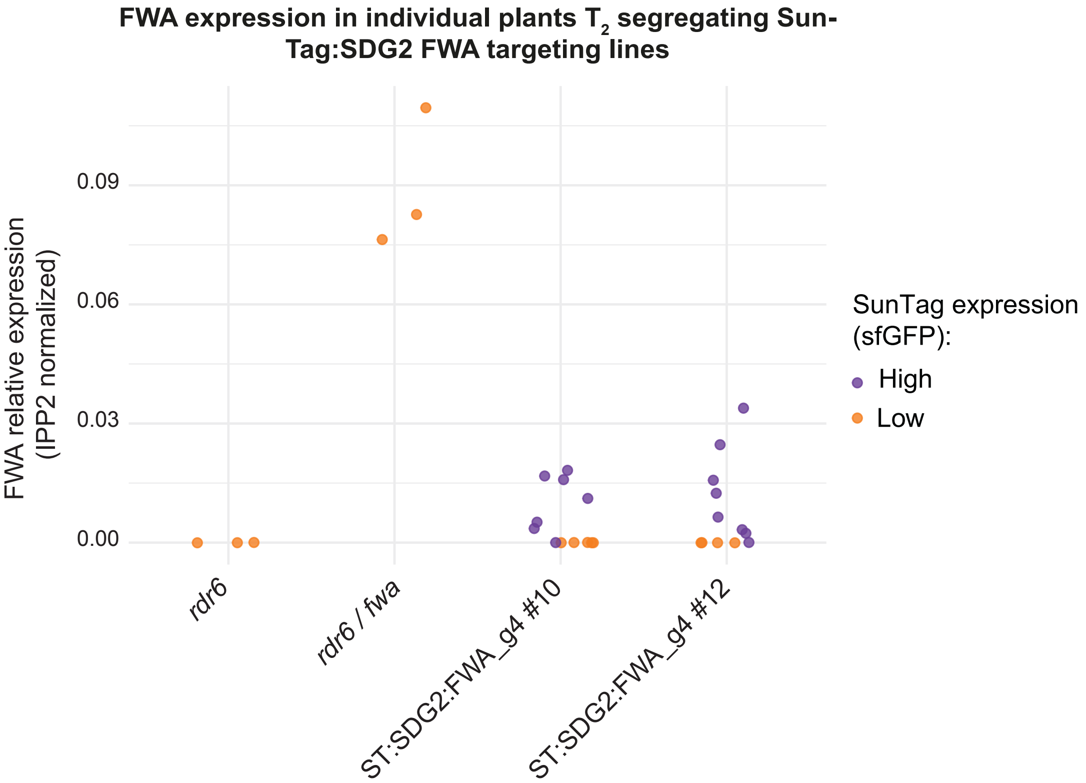

**Supplementary Figure 6:** Transgenerational stability of FWA activation. Plot depicting level of FWA expression (RT-qPCR) in $T_2$ plants from two independent lines segregating for the SunTag:SDG2:FWA_g4 (+/- indicated in purple/orange, respectively). Each dot represents a single $T_2$ plant. n = 3 for *rdr6* and *rdr6 fwa*, n = 12 for *SunTag:SDG2:FWA_g4 #10* and *#12*.

S7

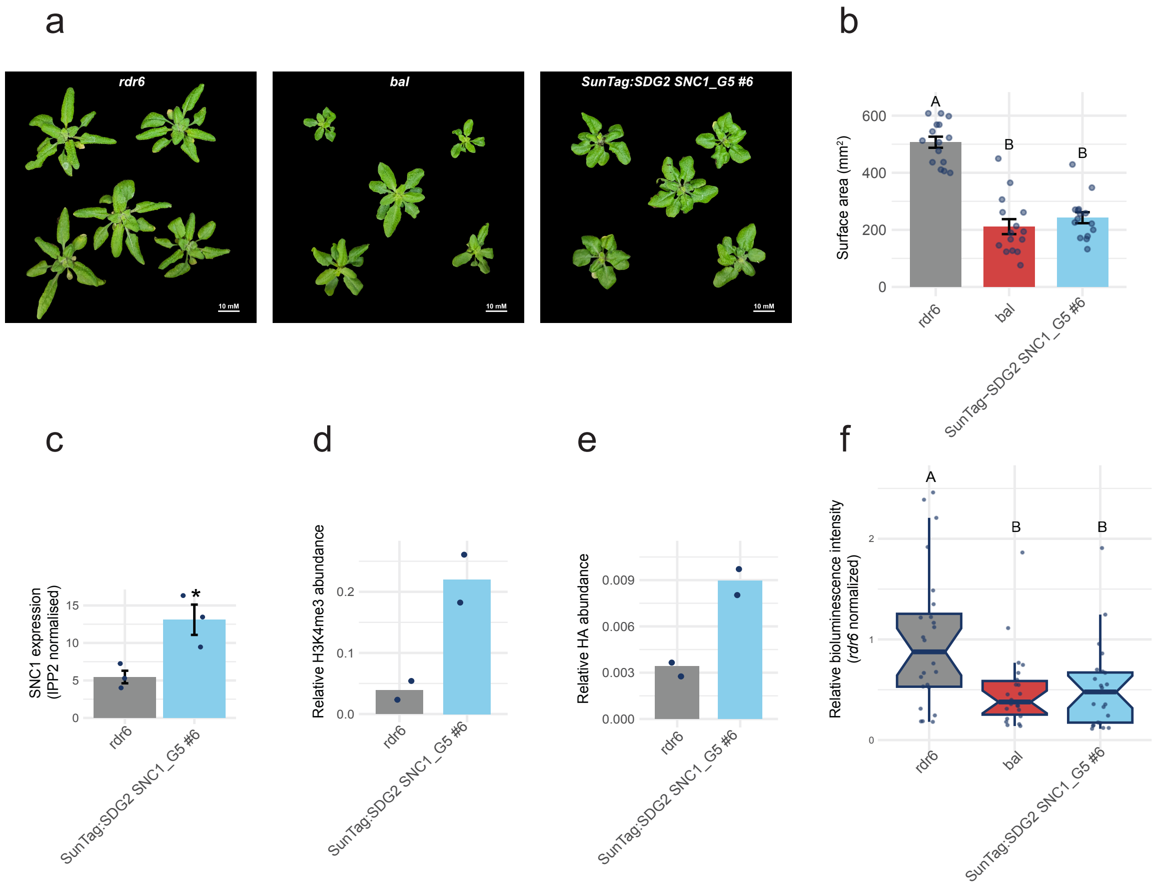

**Supplementary Figure 7:** Independent SunTag:SDG2:SNC1_g5 $T_3$ line targeted to *SNC1* enhances resistance to *P. syringae*. A) representative images of 3-week old plants from the genotypes indicated. Scale bar indicates 1cm. B) Rosette surface area quantification. Each dot represents an individual plant. Error bars represent SEM Different letters indicate significant difference by ANOVA with post-hoc Tukey HSD ($p$ < 0.05). n = 15. C) RT-qPCR for *SNC1* expression. Error bars represent SEM. * indicates p-value < 0.05 evaluated by Student's T-Test. n = 3. D) ChIP qPCR for the presence of SunTag at SNC1. E) ChIP qPCR for H3K4me3 at SNC1. E) *Pst::LUX* assay for colonisation quantification at 3 days post inoculation. Different letters indicate significant difference by ANOVA with post-hoc Tukey HSD ($p$ < 0.05). n = 24.

S8

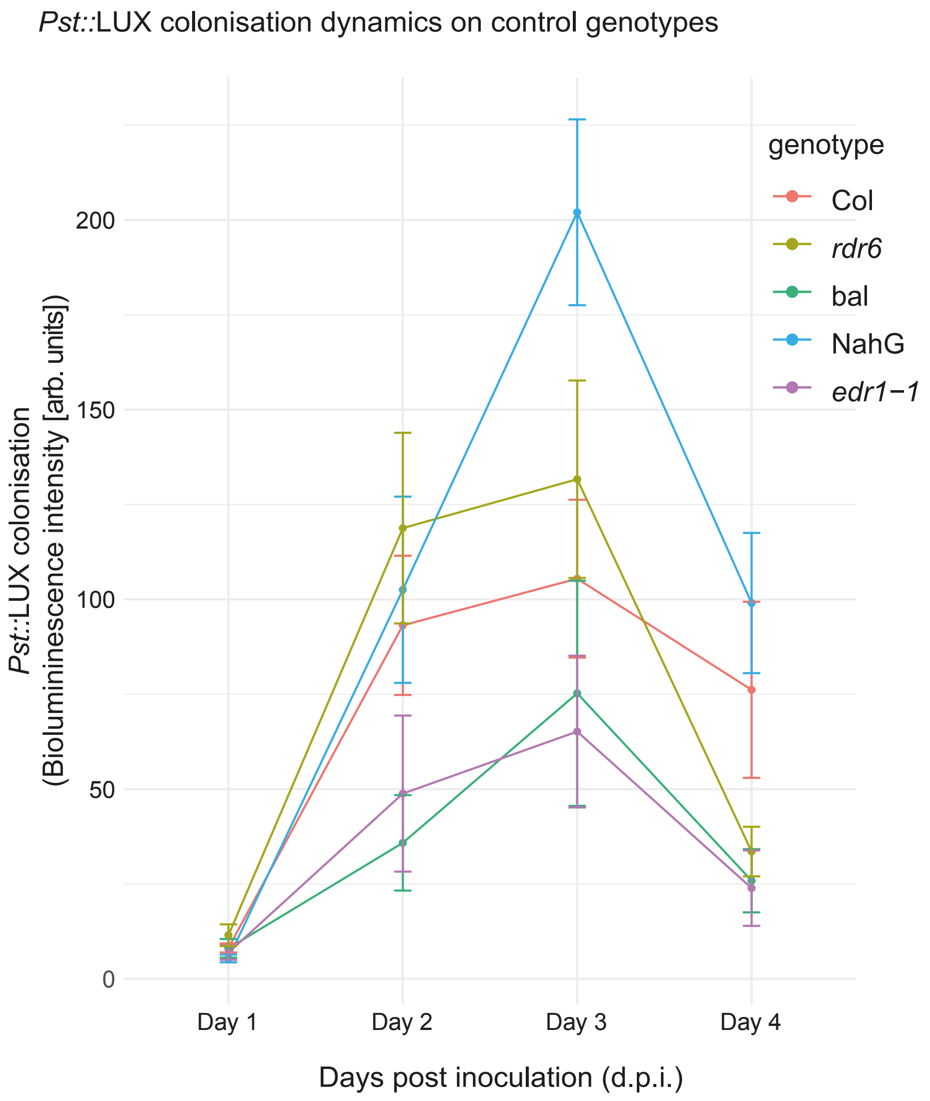

**Supplementary Figure 8:** Colonisation dynamics of *Pst::LUX* in the genotypes indicated. Error bars represent SEM. n = 8 for Col, n = 18 for *rdr6*, n = 10 for *bal*, n = 11 for NahG and n = 7 for *edr1-1*.

S9

**a**

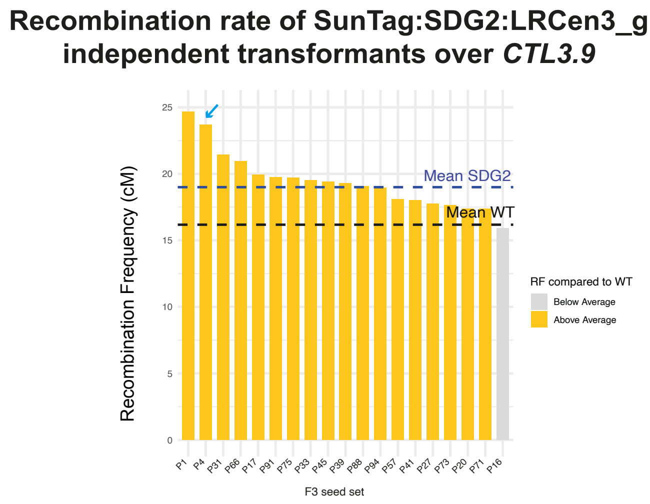
Recombination rate of SunTag:SDG2:LRCen3_g
independent transformants over CTL3.9
Recombination Frequency (cM)
Mean SDG2
Mean WT
RF compared to WT
Below Average
Above Average
F3 seed set
P1 P4 P31 P66 P17 P91 P75 P33 P45 P39 P88 P94 P57 P41 P27 P73 P20 P71 P16

**b**

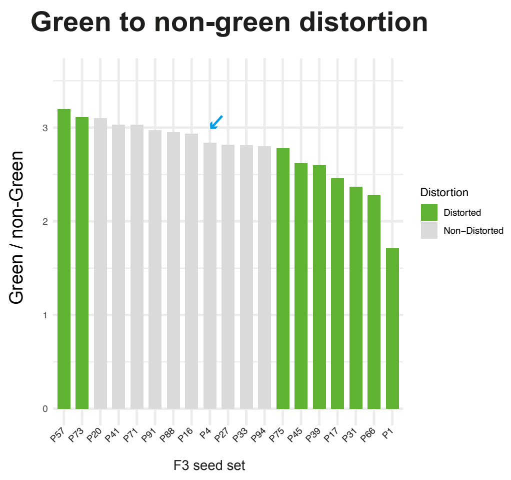
Green to non-green distortion
Green / non-Green
Distortion
Distorted
Non–Distorted
F3 seed set
P57 P73 P20 P41 P71 P91 P88 P16 P4 P27 P33 P94 P75 P45 P39 P17 P31 P66 P1

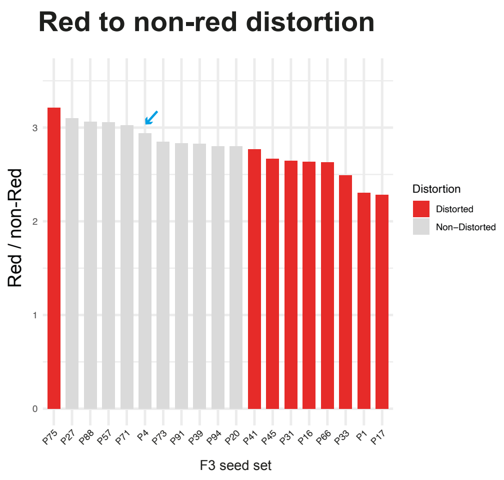
Red to non-red distortion
Red / non-Red
Distortion
Distorted
Non–Distorted
F3 seed set
P75 P27 P88 P57 P71 P4 P73 P91 P39 P94 P20 P41 P45 P31 P16 P66 P33 P1 P17

**c**

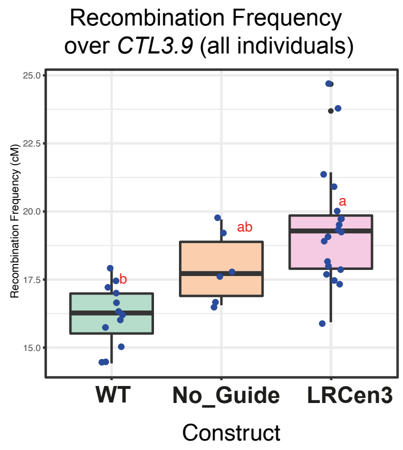
Recombination Frequency
over CTL3.9 (all individuals)
Recombination Frequency (cM)
WT
No_Guide
LRCen3
Construct

**d**

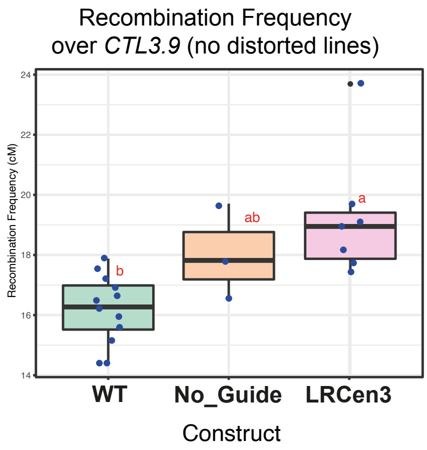
Recombination Frequency
over CTL3.9 (no distorted lines)
Recombination Frequency (cM)
WT
No_Guide
LRCen3
Construct

**Supplementary Figure 9:** A) Recombination rate over *CTL3.9* of the $F_3$ lines (as shown in main Figure 3). Blue arrow (also shown in B) points the P4 line seed set taken to the next generation (also see main Figure 3C, red dot). B) Distortion ratios for the green to non-green (left) and red to non-red (right) T-DNAs from *CTL3.9* from the $F_3$'s. C) Recombination rate over *CTL3.9* ($F_3$'s) including all scorable individuals including no guide lines. n = 12 for control, n = 19 for ST:SDG2-LRCen3 and n = 6 for ST:SDG2-NoGuide. D) Recombination rate over *CTL3.9* ($F_3$'s) filtered for individuals with no distortion (i.e. 3:1). n =12 for control, n = 7 for ST:SDG2-LRCen3 and n = 3 for ST:SDG2-NoGuide. For C) and D) different letters indicate significant difference by ANOVA with post-hoc Tukey HSD ($p<0.05$).

# S10

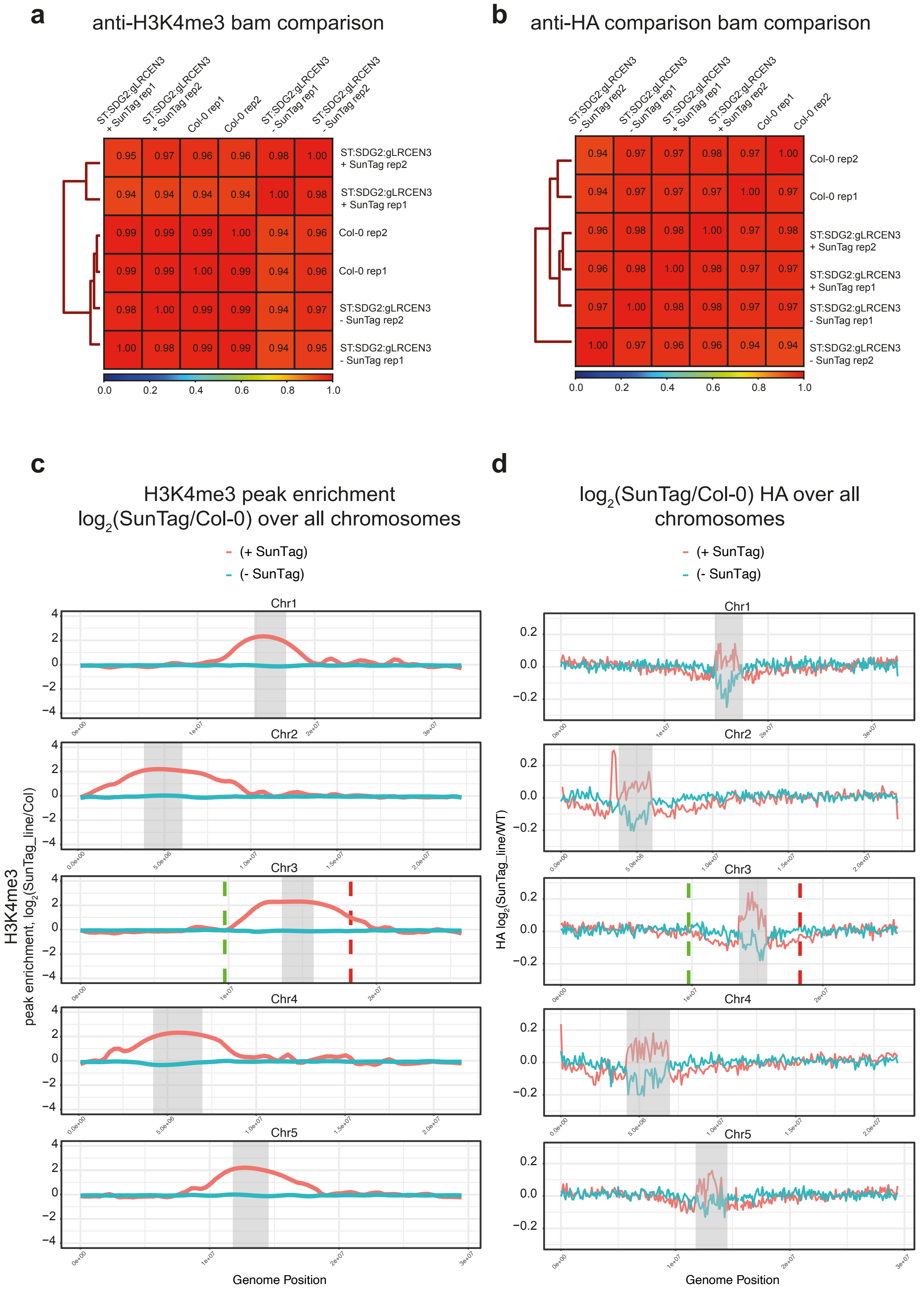

a
anti-H3K4me3 bam comparison
ST:SDG2:gLRCEN3 + SunTag rep1
ST:SDG2:gLRCEN3 + SunTag rep2
Col-0 rep1
Col-0 rep2
ST:SDG2:gLRCEN3 - SunTag rep1
ST:SDG2:gLRCEN3 - SunTag rep2
0.95 0.97 0.96 0.96 0.98 1.00 ST:SDG2:gLRCEN3 + SunTag rep2
0.94 0.94 0.94 0.94 1.00 0.98 ST:SDG2:gLRCEN3 + SunTag rep1
0.99 0.99 0.99 1.00 0.94 0.96 Col-0 rep2
0.99 0.99 1.00 0.99 0.94 0.96 Col-0 rep1
0.98 1.00 0.99 0.99 0.94 0.97 ST:SDG2:gLRCEN3 - SunTag rep2
1.00 0.98 0.99 0.99 0.94 0.95 ST:SDG2:gLRCEN3 - SunTag rep1
0.0 0.2 0.4 0.6 0.8 1.0
b
anti-HA comparison bam comparison
ST:SDG2:gLRCEN3 - SunTag rep2
ST:SDG2:gLRCEN3 - SunTag rep1
ST:SDG2:gLRCEN3 + SunTag rep1
ST:SDG2:gLRCEN3 + SunTag rep2
Col-0 rep1
Col-0 rep2
0.94 0.97 0.97 0.98 0.97 1.00 Col-0 rep2
0.94 0.97 0.97 0.97 1.00 0.97 Col-0 rep1
0.96 0.98 0.98 1.00 0.97 0.98 ST:SDG2:gLRCEN3 + SunTag rep2
0.96 0.98 1.00 0.98 0.97 0.97 ST:SDG2:gLRCEN3 + SunTag rep1
0.97 1.00 0.98 0.98 0.97 0.97 ST:SDG2:gLRCEN3 - SunTag rep1
1.00 0.97 0.96 0.96 0.94 0.94 ST:SDG2:gLRCEN3 - SunTag rep2
0.0 0.2 0.4 0.6 0.8 1.0
c
H3K4me3 peak enrichment
log2(SunTag/Col-0) over all chromosomes
(+ SunTag)
(- SunTag)
Chr1
Chr2
Chr3
Chr4
Chr5
4 2 0 −2 −4
H3K4me3
peak enrichment, log2(SunTag_line/Col)
Genome Position
d
log2(SunTag/Col-0) HA over all chromosomes
(+ SunTag)
(- SunTag)
Chr1
Chr2
Chr3
Chr4
Chr5
0.2 0.0 −0.2
HA log2(SunTag_line/WT)
Genome Position

**Supplementary Figure 10:** A-B) ChIP-seq replicate comparisons by Pearsons correlation clustering using 25bp bin resolution. C-D) ChIP-seq data chromosomal-wide plots as shown in Figure 3, over all chromosomes from sibling lines with or without SunTag:SDG2:LRCen3 (+/-). Enrichment is calculated as $log_2$ fold change over non-transgenic (Col-0) controls in 100kb windows. *CTL3.9* marker positions are indicated, and centromeric regions are shown in grey.

S11

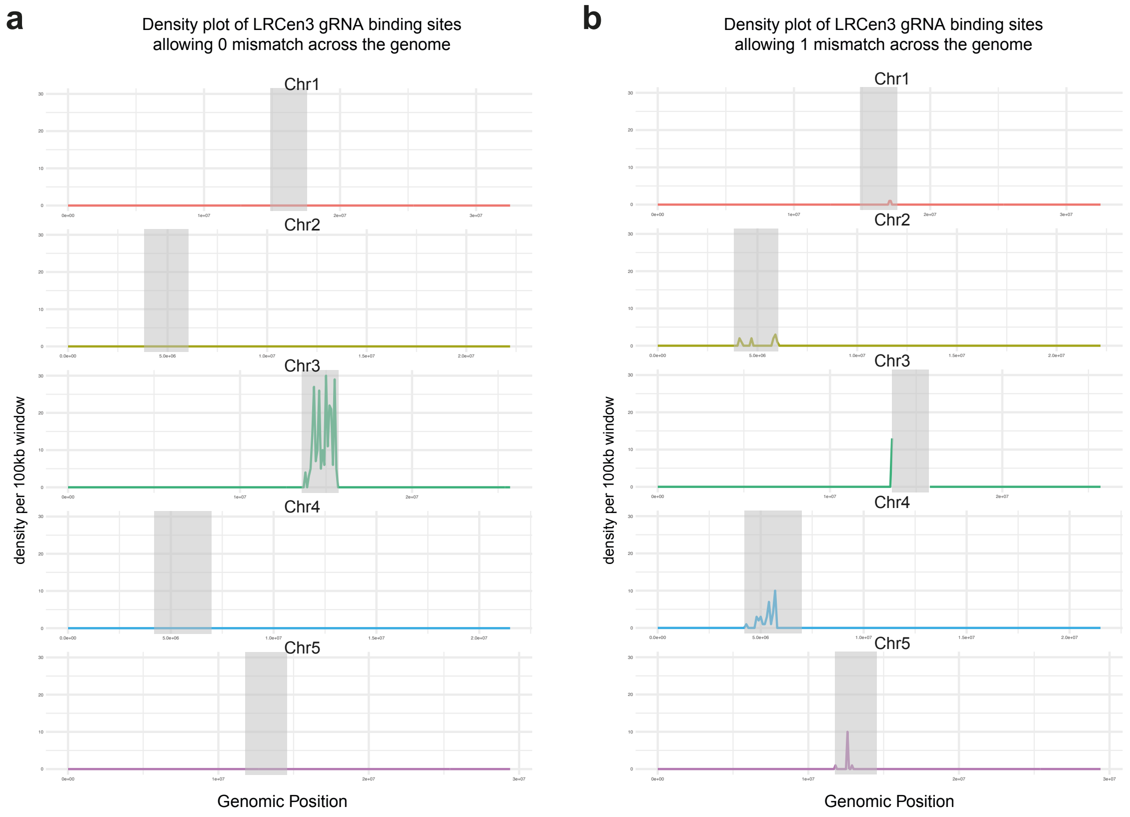

**Supplementary Figure 11:** A) LRCen3 guide RNA binding site density plot with 0 mismatches or B) with 1 mismatch tolerance.

# S12

Centromeric H3K4me3 peaks in SunTag:SDG2:LRCen3 P4 descendents (Col-CEN assembly)

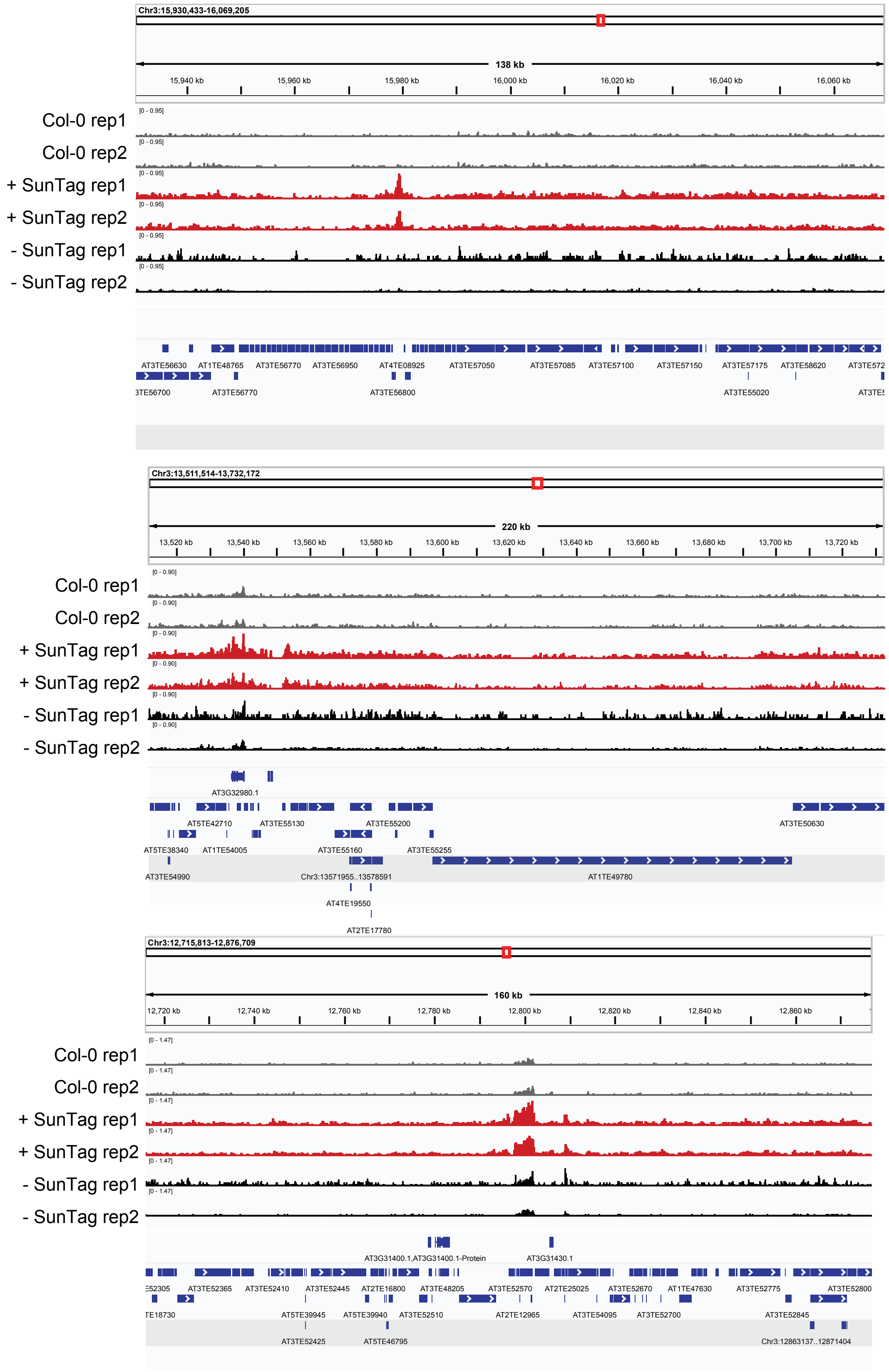

**Supplementary Figure 12:** Representative genome browser images showing examples of centromeric regions where the SunTag:SDG2:LRCen3_g lines show novel or enriched H3K4me3 peaks.

S13

a

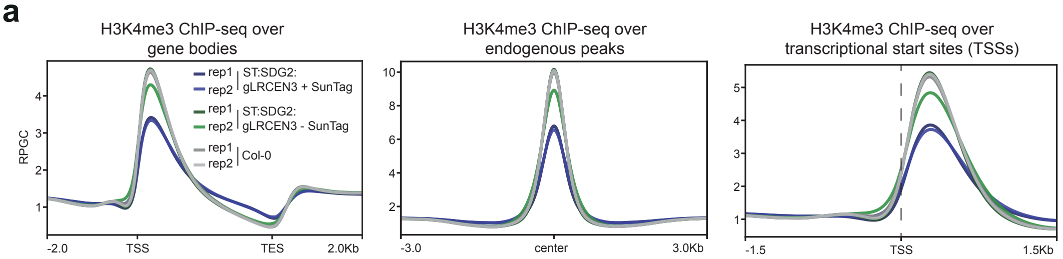

b

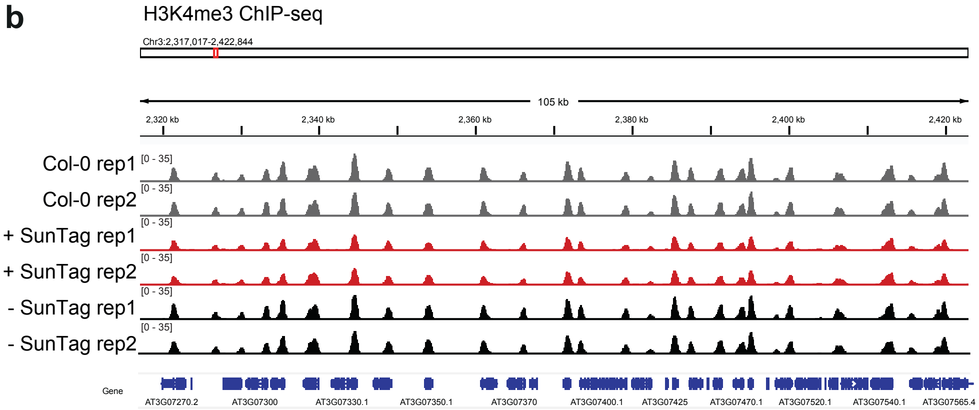

c

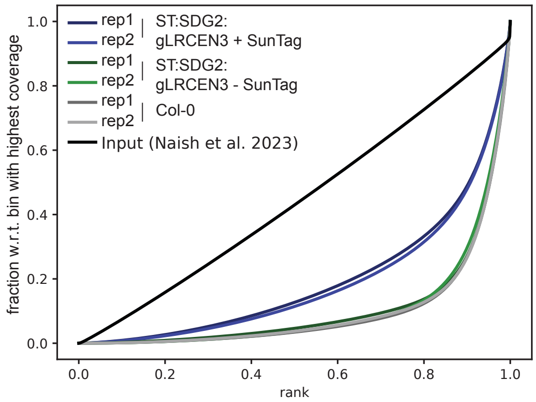

d

| Sample | Total mapped reads | mapped reads in peaks | FrIP (%) |
|---|---|---|---|
| anti-H3K4me3 ST:SDG2:gLRCEN3_withST rep1 | 37124918 | 22335336 | 60.2 |
| anti-H3K4me3 ST:SDG2:gLRCEN3_withST rep2 | 46863315 | 28851483 | 61.6 |
| anti-H3K4me3 ST:SDG2:gLRCEN3_noST rep1 | 50026065 | 40566813 | 81.1 |
| anti-H3K4me3 ST:SDG2:gLRCEN3_noST rep2 | 43236790 | 34269474 | 79.3 |
| anti-H3K4me3 Col0 rep1 | 44335789 | 36150112 | 81.5 |
| anti-H3K4me3 Col0 rep2 | 34744674 | 28141433 | 81.0 |

**Supplementary Figure 13:** Quality control for H3K4me3 ChIPseq of Cen3 targeting SunTag:SDG2 lines. A) Metaplots over genes, H3K4me3 endogenous peaks, TSS regions in the genotypes indicated B) Representative browser image showing the quality of H3K4me3 ChIP-seq data in the genotypes indicated. C) Fingerprint plot (DeepTools ) D) Table of total mapped reads and fragments of reads in peaks (FRiP) scores. Note these are mapped to the TAIR10 assembly (instead of Col-CEN as in Fig. 3 and Supp. Fig. 10-12) to ensure consistency of annotations and for comparison with data shown data from FWA targeting SDG2 and PRDM9 SunTag lines in Fig. 1E and Fig. 4E, Supp. Fig. 5 and Supp. Fig. 20.

S14

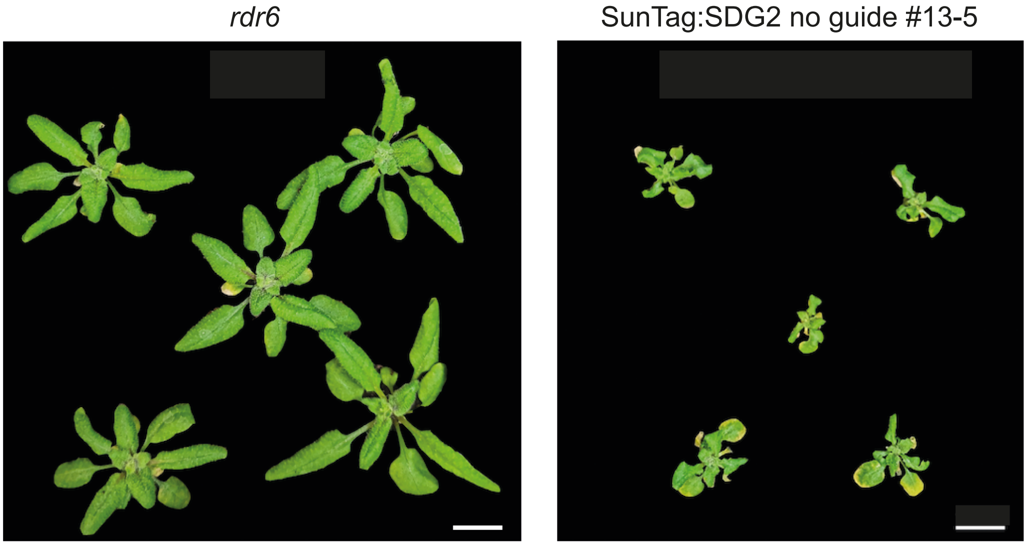

**Supplementary Figure 14:** Dwarfing phenotype of SunTag:SDG2 no guide lines as compared to *rdr6* (untransformed background) controls, representative images from 3-week old plants. Scale bar indicates 1cm.

# S15

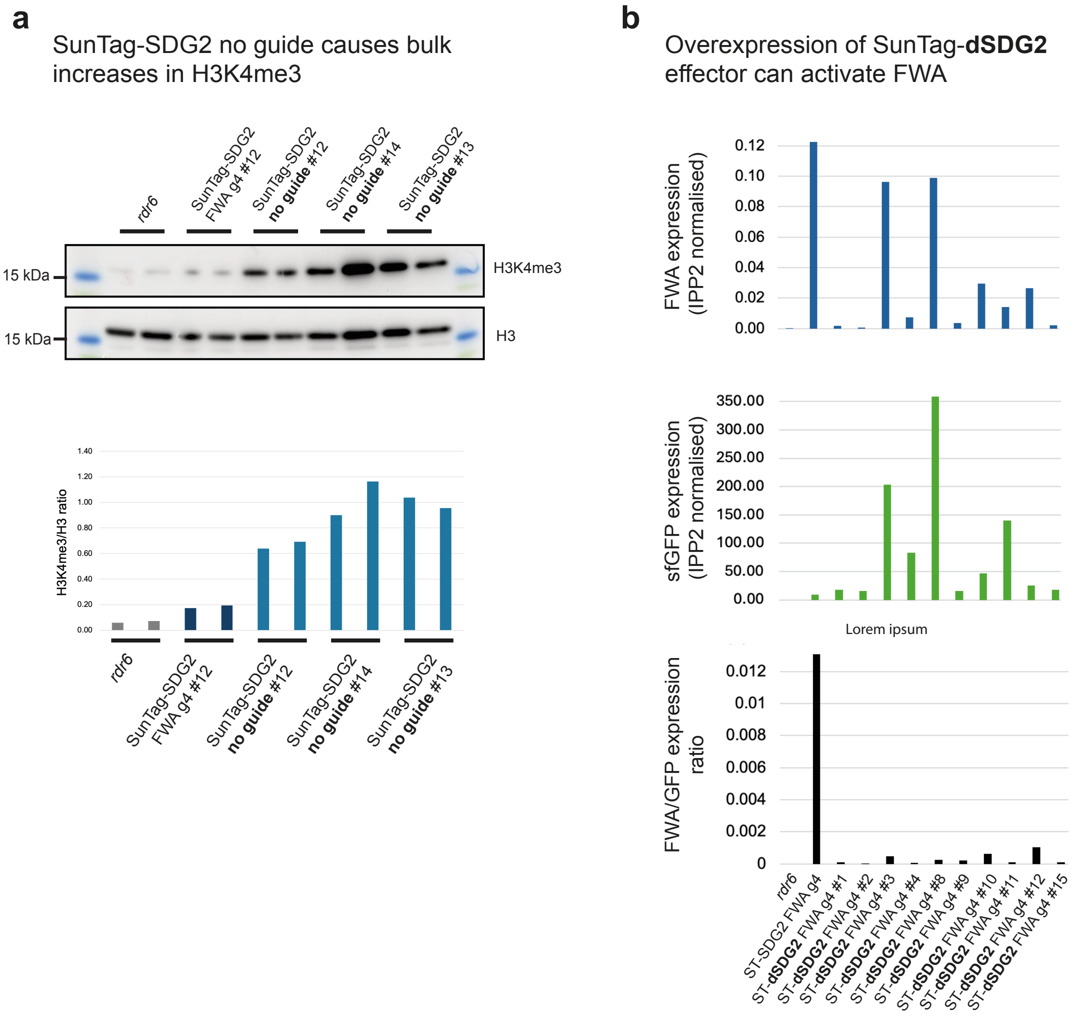

**Supplementary Figure 15:** A) Upper panel, bulk histone western blot showing levels of H3K4me3 in the genotypes indicated, H3 was used as a loading control. Lower panel, quantification of H3K4me3 levels in the genotypes indicated shown as H3K4me3/H3 ratio. B) RT-qPCR for expression of FWA (upper) and sfGFP (middle) and ratio (lower) in the genotypes indicated. Each bar is an individual line, sample is a pool of 10 seedlings.

S16

| Genotype | T1's screened (no.) | FWA expressing (no.) | FWA expressing (%) |
|---|---|---|---|
| SunTag:SDG2:FWA_g4 | 23 | 19 | 83% |
| SunTag:dSDG2:FWA_g4 | 16 | 1 | 6% |
| SunTag:SDG2:No_g | 14 | 2 | 14% |

**Supplementary Figure 16:** Summary table showing the number of FWA activating $T_1$ lines per SunTag:SDG2 construct.

S17

**a**

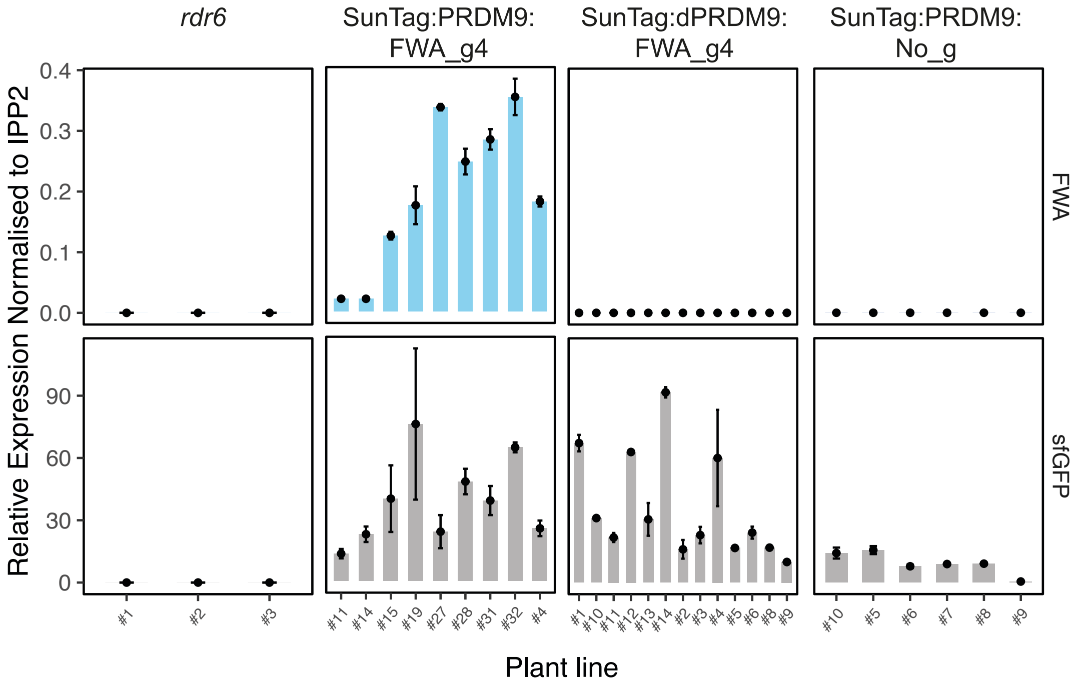

**b**

| Genotype | T1's screened (no.) | FWA expressing (no.) | FWA expressing (%) |
|---|---|---|---|
| SunTag:PRDM9:FWA_g4 | 32 | 23 | 72% |
| SunTag:dPRDM9:FWA_g4 | 19 | 0 | 0% |
| SunTag:PRDM9:No_g | 16 | 0 | 0% |

**Supplementary Figure 17:** A) RT-qPCR of FWA expression (upper) and sfGFP expression (lower) for individual $T_1$ plants from the genotypes indicated. B) Summary table showing the number of FWA activating $T_1$ lines per SunTag:PRDM9 construct.

S18

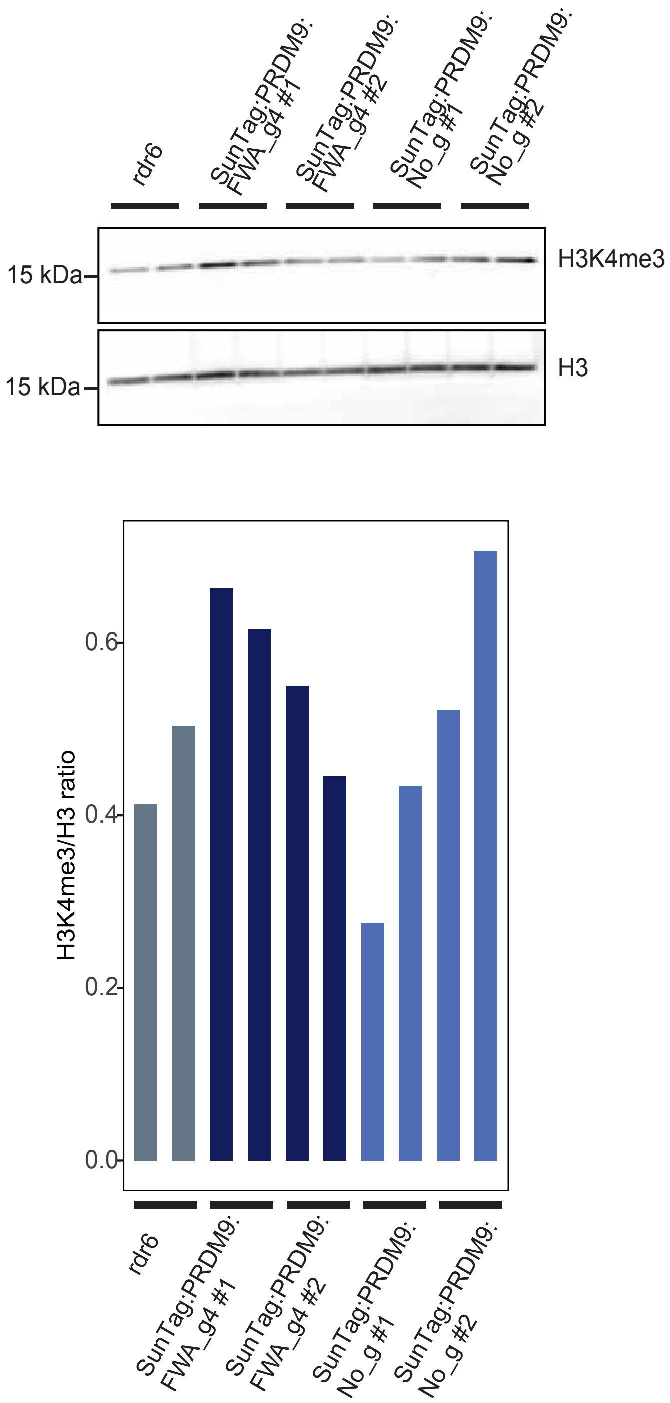

**Supplementary Figure 18:** Upper panel, bulk histone western blot showing levels of H3K4me3 in the genotypes indicated, H3 was used as a loading control. Lower panel, quantification of H3K4me3 levels in the genotypes indicated shown as H3K4me3/H3 ratio.

S19

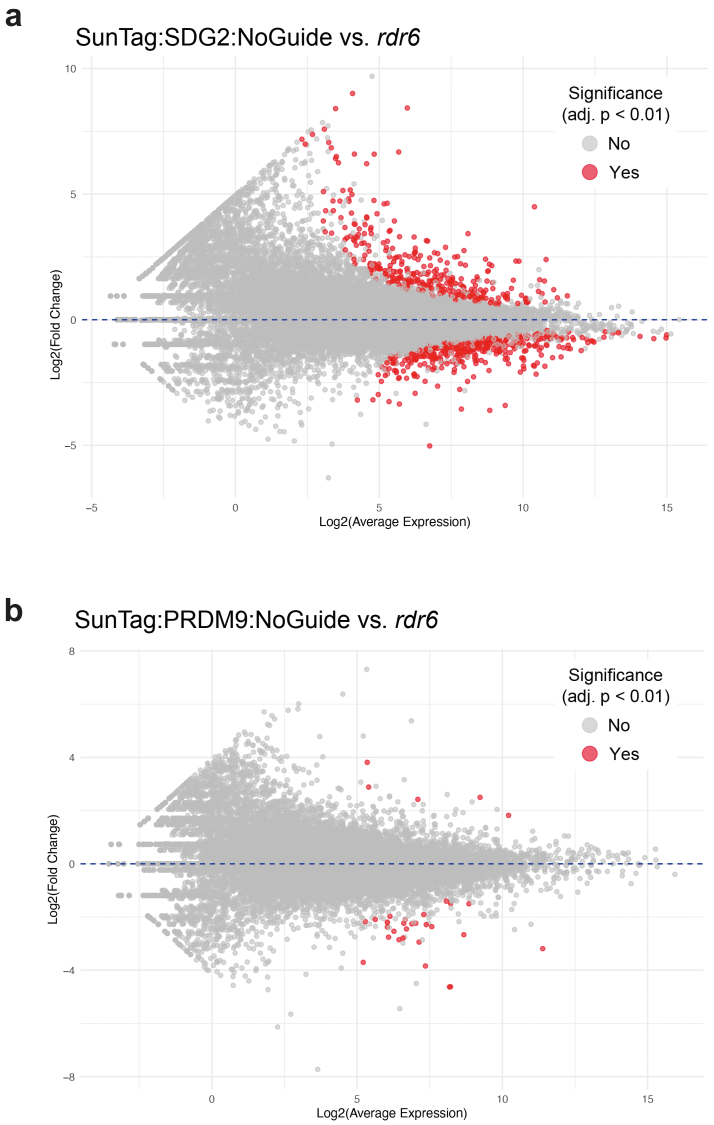

*Note that FWA is not a differentially expressed gene in either comparisons

**Supplementary Figure 19:** A) MA plot for transcriptome of SunTag:SDG2:No_guide versus the non-transformed control (*rdr6*). B) MA plot for transcriptome of SunTag:PDRM9:No_guide versus the non-transformed control (*rdr6*). Differentially expressed genes are depicted in red (FDR < 0.1). Note that FWA is not identified as differentially expressed in either comparison.

S20

**a**

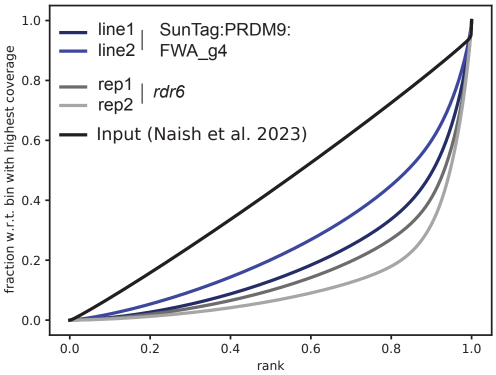

**b**

| Sample | Total mapped reads | mapped reads in peaks | FrIP (%) |
|---|---|---|---|
| SunTag:PRDM9:FWA_g4 line1 | 38519016 | 23382610 | 60.7 |
| SunTag:PRDM9:FWA_g4 line2 | 37332822 | 19114342 | 51.2 |
| rdr6 rep1 | 23207046 | 14856926 | 64.0 |
| rdr6 rep2 | 39630891 | 29971731 | 75.6 |

**Supplementary Figure 20:** Quality control for H3K4me3 ChIPseq of FWA targeting SunTag:PRDM9 lines. A) Fingerprint plot (DeepTools ) B) Table of total mapped reads and fragments of reads in peaks (FRiP) scores.

# S21

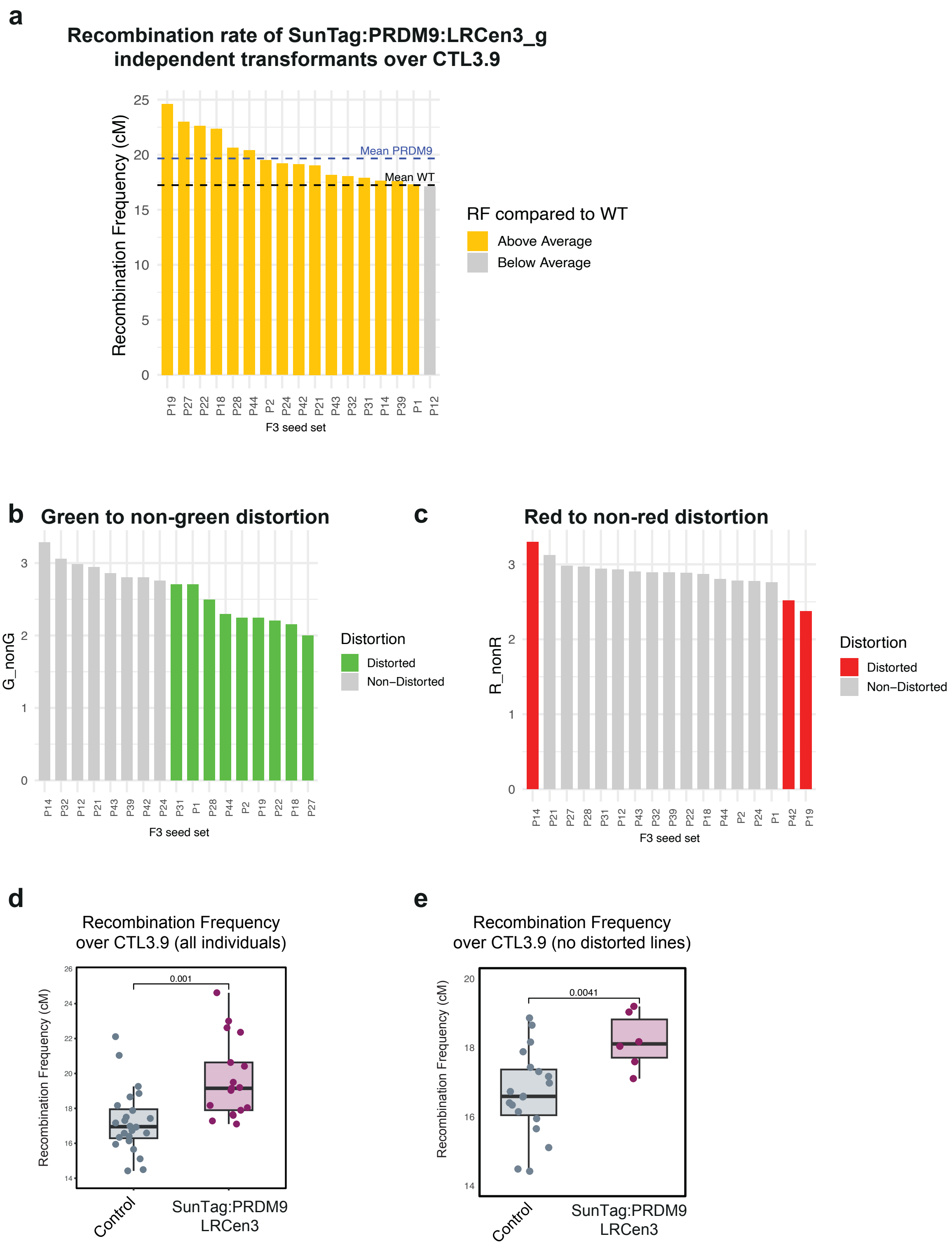
a
Recombination rate of SunTag:PRDM9:LRCen3_g
independent transformants over CTL3.9
Recombination Frequency (cM)
0
5
10
15
20
25
Mean PRDM9
Mean WT
P19 P27 P22 P18 P28 P44 P2 P24 P42 P21 P43 P32 P31 P14 P39 P1 P12
F3 seed set
RF compared to WT
Above Average
Below Average
b
Green to non-green distortion
G_nonG
0
1
2
3
P14 P32 P12 P21 P43 P39 P42 P24 P31 P1 P28 P44 P2 P19 P22 P18 P27
F3 seed set
Distortion
Distorted
Non−Distorted
c
Red to non-red distortion
R_nonR
0
1
2
3
P14 P21 P27 P28 P31 P12 P43 P32 P39 P22 P18 P44 P2 P24 P1 P42 P19
F3 seed set
Distortion
Distorted
Non−Distorted
d
Recombination Frequency
over CTL3.9 (all individuals)
Recombination Frequency (cM)
14
16
18
20
22
24
26
0.001
Control
SunTag:PRDM9
LRCen3
e
Recombination Frequency
over CTL3.9 (no distorted lines)
Recombination Frequency (cM)
14
16
18
20
0.0041
Control
SunTag:PRDM9
LRCen3

**Supplementary Figure 21:** A) Recombination rate over *CTL3.9* of the $F_3$ lines (as shown in main Figure 4). B) Distortion ratios for the green to non-green and C) red to non-red T-DNAs from *CTL3.9* from the $F_3$'s. D) Recombination rate over *CTL3.9* ($F_3$'s) including all scorable individuals. n = 24 for control, n = 17 for Suntag:PRDM9:LRCen3. E) Recombination rate over *CTL3.9* ($F_3$'s) filtered for individuals with no distortion (i.e. 3:1). n = 19 for control, n = 6 for Suntag:PRDM9:LRCen3. *p*-values indicates two-sample students *t*-test.

**Supplementary Table 1:** Oligo List.

| | |
|---|---|
| **SunTag cloning** | |
| SDG2_BsiWI_Fw | GAGGTCGGACCGGTCGTACG |
| SDG2_mutation_Rev | ATTAAAATCGAAAGTTATCTCCTCGCC |
| SDG2_mutation_Fw | GATTTTAATTCTGTAACTGAGAGTAAG |
| SDG2_BsiWI_Rev | CGAGATCCTCCTCCCGTACG |
| Universal F1 | GTCATCTATGTTACGAAGCTTTCGTTGAACAACGG |
| Universal R2 | TCGTGCTCCACCATGGTACCAAAAAAAGCACCGACTCGGTGC |
| pFWA_gRNA4_F2 | acggaaagatgtatgggcttGTTTTAGAGCTAGAAATAGC |
| pFWA_gRNA4_R1 | aagcccatacatctttccgtCAATCACTACTTCGACTCTA |
| pSNC1_gRNA4_F2 | ctatgcaagtcttctaagaaGTTTTAGAGCTAGAAATAGC |
| pSNC1_gRNA4_R1 | ttcttagaagacttgcatagCAATCACTACTTCGACTCTA |
| pSNC1_gRNA5_F2 | atcaaatccctaacaagataGTTTTAGAGCTAGAAATAGC |
| pSNC1_gRNA5_R1 | tatcttgttagggatttgatCAATCACTACTTCGACTCTA |
| LRCen3_g_F2 | aggcttacaagattgggttg**GTTTTAGAGCTAGAAATAGC** |
| LRCen3_g_R1 | caacccaatcttgtaagcct**CAATCACTACTTCGACTCTA** |
| **PRDM9 cloning into SunTag** | |
| RP_PRDM9_G282A_3' | cgagatcctcctcccgtacgtcaaa |
| FP_PRDM9_G282A_5' | GAGGTCGGACCGGTCGTACGTATC |
| RP_PRDM9_G282A_middle | CTCTGTGATCTGAGCCTCATAGGGGCCA |
| FP_PRDM9_G282A_middle | TGGCCCCTATGAGGCTCAGATCACAGAG |
| **ChIP-qPCRs** | |
| High_H3K4me3_AT5G65130_F | CCAGACATTGTTCGTCAAGGACAC |
| High_H3K4me3_AT5G65130_R | CAACGTTATATCCGACTCAGGAGATG |
| Low_H3K4me3_AT2G40004up_F | CAGCTACAAGTTCAACTAATCATGC |
| Low_H3K4me3_AT2G40004up_R | CTTTAATTGCCCGTCATTACGCTAC |
| pFWA_ChIP_Fw | AATATCAGATCTTGCGCCGCTC |
| pFWA_ChIP_Rev | GAAAAACAGACAAATCGGGAACC |
| pSNC1_ChIP_Fw1 | ATGGCCTTATCTTGTTAGG |
| pSNC1_ChIP_Fw2 | GAGTAGAAGCTTTCCAAGG |
| **RT-qPCRs** | |
| sfGFP_qPCR_Fw | CAAAGATGACGGGACCTACAA |

| | |
|---|---|
| sfGFP_qPCR_Rev | GTACTCGAGTTTGTGTCCAAGA |
| FWA_qPCR_Fw | TTAGATCCAAAGGAGTATCAAAG |
| FWA_qPCR_Rev | CTTTGGTACCAGCGGAGA |
| SNC1_qPCR_Fw | TTTGAAGAGACATGCAAGGC |
| SNC1_qPCR_Rev | TCGGCAAGCTCTTCAATCATGG |
